# Genomic diversity analysis enables development of pan-Dengue Toehold RNA sensors

**DOI:** 10.1101/2025.08.23.671906

**Authors:** Anirudh Nandakumar, Deeksha Mishra, Akshay Shendre, Edrea Mendonca, Akash Gulyani, Arati Ramesh

**Author notes:** contributed equally to this work (Anirudh Nandakumar, Deeksha Mishra). co-corresponding authors: Correspondence should be addressed to: Arati Ramesh, Tata Institute for Genetics and Society, GKVK Campus, Bellary Road, Bangalore, 560065,; Akash Gulyani, Department of Biochemistry, School of Life Sciences, University of Hyderabad, Hyderabad, India.

## Abstract

The Dengue virus (DENV) like many other RNA viruses exhibits high genome sequence diversity. This poses a challenge to nucleic acid-based diagnostics, rendering them inefficient at detecting the diverse DENV strains circulating in a population. In this study, we address this challenge by developing a Toehold sensor assay that despite significant genomic diversity is able to detect ∼99.4% of all strains of DENV. To this end, our workflow first identifies relatively conserved short stretches (36-nt) within all DENV genomes, which could potentially serve as triggers that activate Toehold RNA sensors. We then add a crucial step of mismatch-tolerance that allows related triggers with high sequence diversity and multiple mismatches in the sensor-binding region to be efficiently sensed by the same sensor. The sensitivity of the Toehold sensor assay is often increased by inclusion of an isothermal RNA amplification step. We show by employing multiple primer sets that diverse trigger RNAs from all four serotypes of the dengue virus can be successfully amplified and subsequently detected by the same sensor. Together, this approach has resulted in a *pan-dengue* Toehold sensor assay, which presents a powerful nucleic acid detection platform to detect viruses with high sequence diversity.

## INTRODUCTION

Over 300 different viruses are known to infect humans and cause disease. Timely detection of infection is important not only to guide treatment but also to prevent the spread of infection. One factor that continues to obscure detection is the enormous sequence diversity observed in viral genomes. The genome sequence diversity observed in viruses can be attributed to multiple factors such as the inherent rate of mutation, recombination, reassortment and antibody-mediated selection pressure ^1–4^. DNA viruses have mutation rates between 10^−8^ to 10^−6^ substitutions per nucleotide site, per cell infection and this is even higher in RNA viruses which show mutation rates of 10^−6^ to 10^−4^ substitutions per nucleotide site, per cell infection^5,6^.

Dengue (DENV), a positive-strand RNA virus, shows high sequence diversity, exemplified by the fact that it exists as at least four distinct serotypes 1, 2, 3, and 4^7–9^, which themselves are sub-divided into multiple genotypes. The Dengue virus causes significant morbidity, posing a major burden to healthcare systems particularly in tropical and sub-tropical countries where dengue is widely prevalent. Dengue infection may cause symptoms ranging from mild fever, severe flu like symptoms, to hemorrhagic fever including serious bleeding, drop in blood pressure and death. As per WHO estimates, there are about 100 to 400 million dengue infections per year, globally. Estimates suggest that as many as 3.9 billion people are at the risk of dengue infection ^10,11^. This makes dengue detection a pressing challenge, requiring innovative approaches.

Diagnosis of dengue infection is often done through immunoassays against the NS1 protein^12,13^, whose signature regions also help to type the virus^14^. Alternately, detection of the viral RNA through RT-qPCR assays is a recommended diagnostic test especially in the acute phase of infection^15^. Several nucleic acid detection strategies such as NASBA^16–18^, RT-RPA^19–21^, CRISPR-based diagnostics ^22,23^, and RT-LAMP^24,25^ have been described, where amplification of viral RNA relies on sequence-dependent primer/guide RNA recognition. Notably, in all these methods, the sequence diversity inherent to the dengue genomes could obscure diagnosis.

To address viral sequence diversity, one approach has been to align representative DENV sequences from all genotypes/serotypes to identify the most conserved segments of a specific genomic region such as the C-prM and NS1 genes, or the 3’ UTR of the genome^25–28^. Other approaches involve exploiting the mismatch tolerance of primer-target recognition^24^, or engineering mismatch tolerance into the amplification reaction by adding a proof-reading polymerase which removes 3’ end mismatches^29^. These studies along with recently reported tools for analyzing viral sequence diversity^30,31^ underline the importance of considering sequence diversity while developing assays for viral detection.

A recent innovation in viral detection are the toehold RNA sensors, which have been designed to detect viruses including Zika^32^, SARS-CoV-2^33,34^, hepatitis A^35^ and norovirus^36^. Toehold sensors have also been adapted to detect pathologically relevant miRNA^37^ and for profiling common gut-bacteria^38^. The toehold RNA switch is a structured element placed upstream of a reporter mRNA, whose ribosome binding site (RBS) and start codon (AUG) are sequestered in a base-paired structure and hence kept inaccessible to the ribosome for translation. The sensors are designed to contain a sequence complementary to the target, that upon binding the target RNA, initiates a structural rearrangement of the sensor, enabling translation of the reporter gene. The sensors can be coupled to various reporters that enable visual (lacZ), portable (gfp, glucose oxidase), and rapid (nano-lantern) detection of the target^32,33,39,40^. When coupled with nucleic acid amplification techniques such as NASBA^32,33,36^, RT-RPA^36^, or RT-LAMP^34^ these sensors show sensitivity comparable to RT-LAMP and RT-RPA assays. While these studies reveal that toehold RNA-based detection is versatile and meets the needs of diverse diagnostic settings, they do not comprehensively address the sequence diversity of the viral target itself. One example of viruses that are highly divergent in sequence is the norovirus. To design toehold sensors for noroviruses, a conserved region was found using sequence alignments of multiple norovirus genomes and toehold sensors were designed against the conserved region^36^. Since viruses continue to accumulate mutations over time, it becomes essential to employ a dedicated design strategy that takes into consideration this sequence divergence.

In this work, we have designed and developed a toehold RNA-based assay that overcomes DENV sequence diversity and results in a *pan-dengue* assay. For this, we established a computational pipeline that first identifies the 36-nucleotide regions with maximum conservation among all DENV genomes. We then exploit the inherent ability of a toehold sensor to tolerate a certain degree of mismatch with its target sequence, resulting in related triggers from very divergent DENV genomes to be sensed by the same toehold sensor. This strategy resulted in DENV-toehold sensors that could detect >99.4% of DENV sequence variations globally reported thus far, making them truly “*pan-dengue*” sensors. Typically, toehold sensor assays are coupled with an RNA amplification step for increased sensitivity of detection. Here, using multiple primer sets we show that diverse RNA triggers belonging to each of the four DENV serotypes could be successfully amplified and sensed by our *pan-dengue* sensor.

## RESULTS

### Analysis of viral genomes for sequence level diversity

We analyzed all the sequences of *Betacoronavirus cameli* (MERS), *Orthoflavivirus zikaense* (ZIKV), *Orthoflavivirus denguei* (DENV), and *Norovirus norwalkense* (NORO), to estimate the diversity in the nucleotide sequence among their genomes. Full-length genome sequences of MERS (219 genomes), ZIKV (729 genomes), DENV (6712 genomes) and NORO (1452 genomes) were taken from the Bacterial and Viral Bioinformatics Resource Center^41^ database, and the Shannon entropy^42,43^ for every position across the genome was calculated (Figure 1A-B). DENV and NORO genomes show high divergence at several genomic positions (Shannon entropy ≥ 1 in 23% and 27% of total positions respectively). In contrast, MERS and ZIKV contain no positions with Shannon entropy ≥ 1).

**Figure 1:**
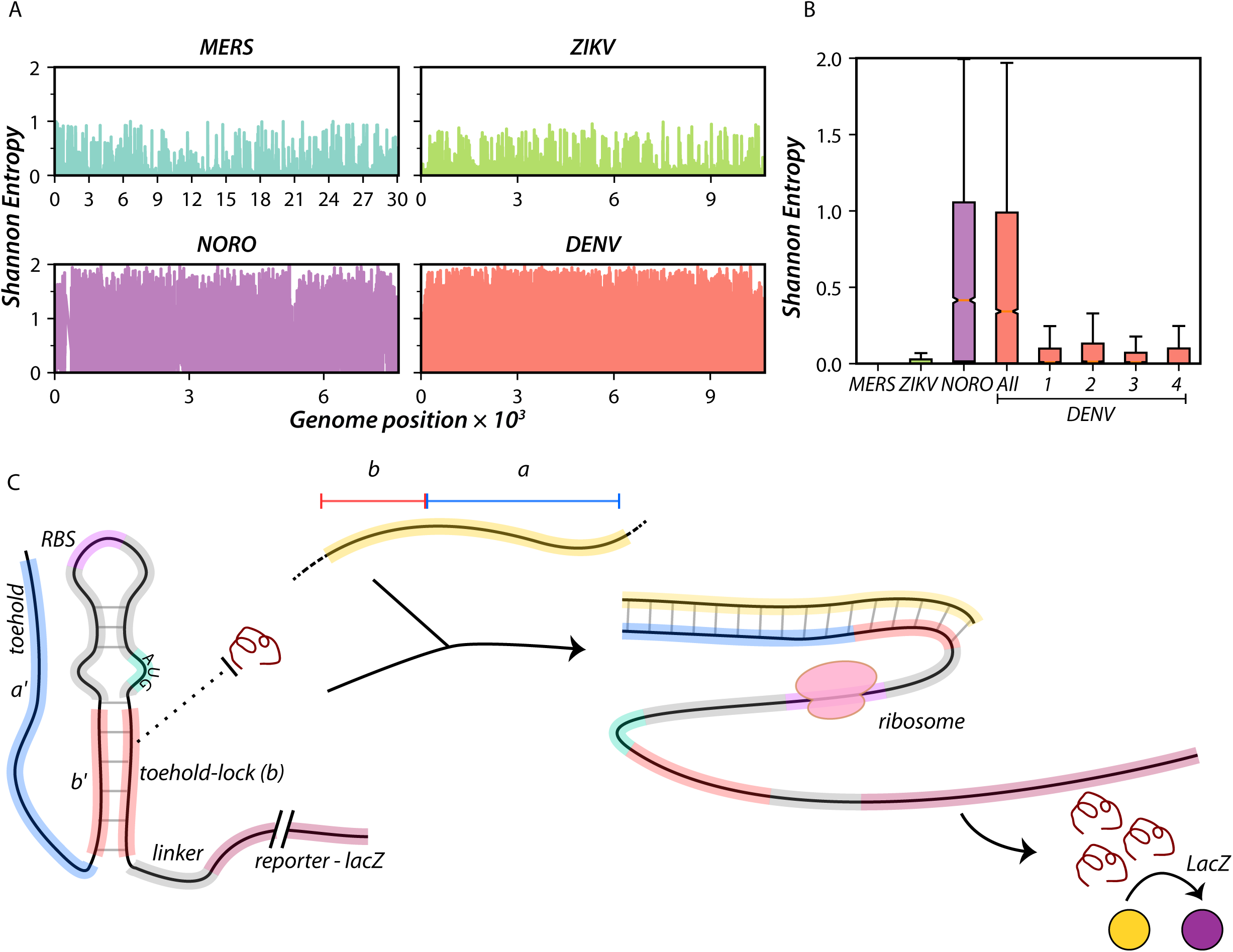
Nucleotide-level diversity in RNA viruses and concept of Toehold sensors. A) Shannon entropy plots at each nucleotide position across the genomes of four ssRNA viruses are shown. MERS (tax. id: 1335626, 219 genomes), ZIKV (tax. id: 64320, 729 genomes), NORO (tax. id: 142786, 1452 genomes), and DENV (tax. id: 12637, 6826 genomes) are shown. Shannon entropy values =2 indicates no conservation. B) Average Shannon Entropy calculated across every nucleotide position is shown for MERS, ZIKV, NORO, DENV, and four DENV serotypes. Bars represent the mean, and error bars represent standard deviation. DENV and NORO exhibit high sequence diversity throughout the genome, compared to MERS and ZIKV. Each serotype of DENV also exhibits higher levels of diversity within them compared to MERS and ZIKV. C) The toehold RNA sensor in its off state, is composed of a structured region (gray and red stems) juxtaposing the RBS (purple) and the start codon AUG (cyan). This structure impedes translation initiation of the downstream reporter (such as *lacZ*). The trigger RNA (yellow) binds the toehold region (blue) and extends into the toehold-lock, thus unzipping it. This allows access of the ribosome to the RBS and start codon, upregulating translation of the reporter protein.

Dengue is caused by four different serotypes of DENV, each with a distinct serum neutralization profile. The Shannon entropy calculations reveal a high level of sequence divergence even within each serotype. While the mean Shannon Entropy for DENV and NORO are 0.539 and 0.594, and for MERS and ZIKV are 0.007 and 0.026, the mean Shannon Entropy for the individual DENV serotypes are in the range of 0.117 to 0.164. (Fig. 1B and Fig S1).

This degree of sequence divergence implies that identification of conserved genomic regions in viruses like DENV and NORO is challenging. Any *pan-dengue* biosensors designed to report on all dengue variants would ideally need to take into consideration this diversity. We decided to address this diversity in our design of the toehold-RNA based dengue sensors.

### Design of toehold biosensors for detection of Dengue

The toehold sensor contains a 36-nucleotide region that is complementary to a portion of the target viral RNA (Fig 1C). 11 nucleotides at the 3’ end of the 36-nt stretch are part of a base-paired structure that keeps the RBS and AUG sequestered, while the initial 25 nucleotides toehold region that engages the target (trigger) RNA is linear. Trigger-binding initiates the disruption of the structure, allowing the translation of the downstream reporter (Fig 1C).

To identify 36-nt trigger sequences that are conserved across all DENV serotypes, we took all full-length DENV genomes (6712 genomes) from the BV-BRC database and indexed all possible 36-nucleotide triggers in *each* genome. Occurrence of each of these triggers across every DENV genome was calculated. We found that even the relatively more conserved triggers appeared in 97% and 72% of genomes of a serotype but were absent in other serotypes (Fig. 2A left panel).

**Figure 2).**
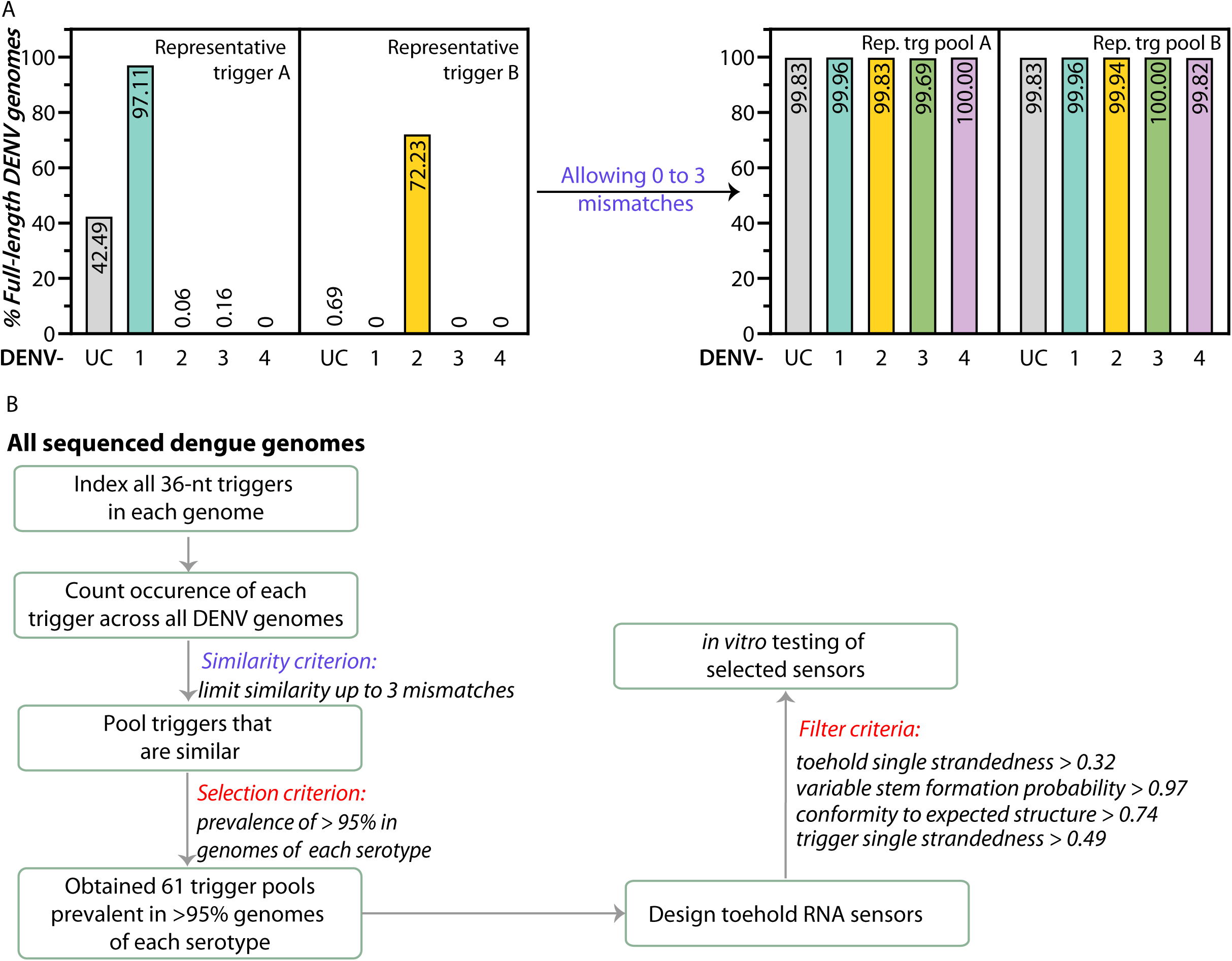
Design of Toehold RNA sensors to detect all variants of Dengue virus. A) Left: Percentage of full-length DENV genomes in which a representative trigger is found, in each serotype DENV 1-4 and unclassified (UC). Pooling triggers that have 0 to 3 mismatches with Trigger A and B leads to trigger pools that are present in >99.6% of DENV genomes in each serotype (Right). (B) Potential pan-Dengue toehold sensors were designed using a computational pipeline outlined in the flowchart. All available high-quality full-length genomes of DENV from DENV 1,2,3,4 and UC were taken. A library of unique 36-nt triggers was built and occurrence across all DENV genomes was calculated. To address low occurrence of triggers across genomes, a *Similarity Criterion* was applied where every trigger was pooled with triggers that varied by 0 to 3 mismatches (Hamming distance). A *selection Criterion* was applied where only trigger pools that were present in >96% genomes across each serotype were retained. Toehold sensors were designed to detect the parent trigger of each trigger pool. The sensors and their cognate triggers were filtered on the basis of four secondary structure parameters indicated, and taken forward for *in vitro* testing.

We then added a crucial step that allows mismatches between the Trigger and sensor region. This relaxation of trigger conservation was inspired by previous studies^36^ which have shown that toehold sensors can also be turned on by RNA triggers that have up to 3 mismatches. Hence, we took triggers which had 0, 1, 2, or 3 sequence mismatches (within 3 Hamming distances) and pooled them together. The resulting trigger pools were selected if they were prevalent in >96% of genomes of each serotype. For example, applying the similarity criteria to the two representative triggers shown in Fig 2A results in trigger pools that appear in >99.6% of genomes of each serotype (Fig 2A right panel, Table S1). Totally, we obtained 61 trigger pools that appear in >96% of genomes of each serotype.

Interestingly, all the 61 trigger pools map to an open reading frame encoding the capsid protein close to the start of the ORF (Fig S2). Thus, our analysis suggests that the region within the sequence coding for the capsid protein is the least divergent in terms of sequence. The 3’ UTR of the DENV genomic RNA has been shown to make long range interactions with the 5’ of the genome and these interactions are important for replication of the DENV genome^44,45^. The region to which our trigger pools map, is included in this previously noted long range interaction.

These 61 trigger pools were used to design the corresponding toehold sensors (Fig. 2B and Methods). We scored the triggers and sensors on four criteria such as the single strandedness of the trigger, toehold single strandedness, probability of lower stem formation, and conformity to the designed structure of the sensor were calculated for all the designed sensors. We noted the average trigger and toehold single-strandedness of the targets and sensors were 0.54 and 0.46 (Fig 3A), respectively. These values are lower than those used for selecting toehold sensors designed against SARS-CoV-2^33^ and suggest a higher degree of intra-molecular interactions in the DENV target RNAs and concomitantly in the toehold region of DENV sensors.

**Figure 3:**
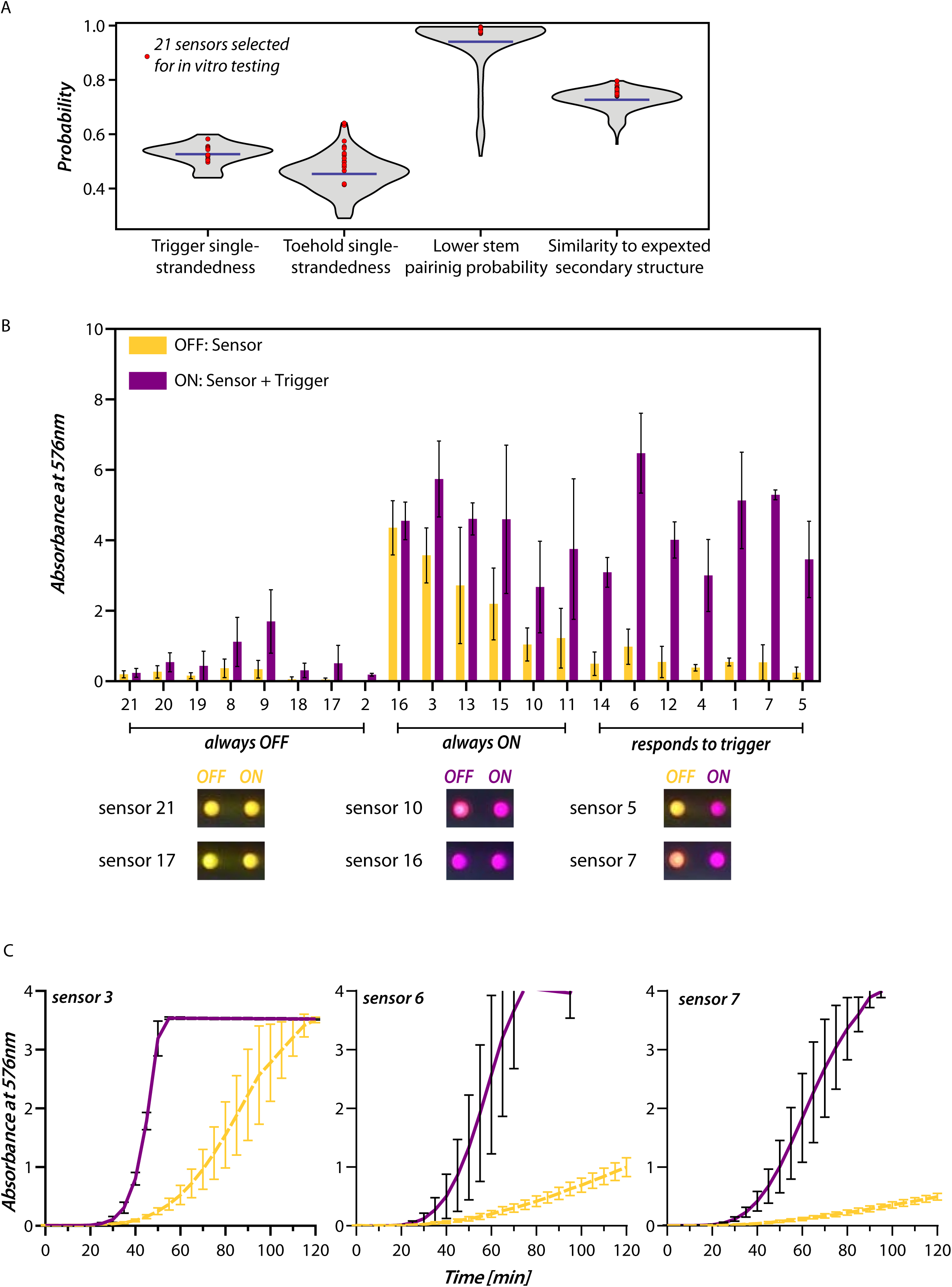
*In vitro* screening of DENV Toehold RNA sensors. A) Violin plot (gray) shows the distribution of the four secondary structure parameters of 61 triggers and 219 toehold sensors. Mean value of the parameters is indicated by the blue line. Parameters of the 21 sensors chosen for in vitro testing are shown (Red circle). B) In vitro transcription translation (IVTT) assay of 21 sensors in presence (purple) or absence (yellow) of their cognate triggers is shown. Absorption of CPRG dye at 576 nm was used to report the extent of translation at the end of 2h at 37℃. The sensors showed three types of responses-those that did not turn ON even in the presence of the trigger (always OFF), those that were ON even in the absence of trigger (always ON), and sensors that responded specifically to the trigger. Post-2hour images of the IVTT reaction are shown for select sensors from the three categories mentioned above (also see Fig S3). Bars represent mean and the error bars represent standard deviation (n=3). C) Absorbance at 576 nm was monitored over time for IVTT reactions with sensors 3, 6, and 7. Presence of trigger (purple) showed faster increase in absorbance when compared to absence of trigger (yellow). Lines represent mean and the error bars represent standard deviation (n=3).

### Toehold sensors that report efficiently on the presence of dengue RNA

We tested 21 candidate toehold sensors coupled to the *lacZ* reporter. We measured their response to their corresponding trigger in terms of the increase in absorbance at 576 nm due to the hydrolysis of CPRG by LacZ at the end of two hours (Fig. 3B, S3A). DNA corresponding to each candidate toehold sensor was used as input for an *in vitro* transcription translation (IVTT) coupled assay. *In vitro* synthesized trigger RNAs were used to check the performance of the sensors. The IVTT reactions showed three types of responses. Sensors (21, 20, 19, 8, 9, 18, 17, and 2) didn’t turn ON even in the presence of their triggers (ON < 2), sensors (16, 3, 13, 15, 10, and 11) were constitutively turned ON independent of the trigger (ON/OFF ≤ 3), and sensors (14, 6, 12, 4, 1, 7, and 5) showed a discernible change in the presence of trigger RNA compared to its absence (ON/OFF > 5). We chose sensors 3, 6, and 7 with the highest ON states, and performed time course experiments (Fig 3C). We noticed that sensor 3 had a high ON/OFF ratio in the 30 to 40-minute time range and saturates faster (A_576nm_ = 4 in 30 minutes) than sensor 6 (65 minutes) and sensor 7 (115 minutes).

### Validation of sensors for their ability to detect all Dengue serotypes

We next asked if our sensors would be capable of detecting the majority of DENV genome variants. This is important given that our *pan-dengue* sensors could only be designed when we made an allowance of 0 to 3 mismatches while identifying triggers that were present in more than 95% genomes of each serotype. To test the mismatch tolerance of our sensors, we synthesized several triggers from the trigger pools for sensors 3, 6, and 7, and tested them in an IVTT assay. For sensor 3, we tested 13 different RNA triggers that represented 99.6% of all DENV genomes (Fig. 4A and Fig S4). Sensor 3 shows a good response to all tested triggers including those with 3 mismatches between the trigger and the toehold region. Similarly, for sensor 6 (Fig 4B) and sensor 7 (Fig 4C), we tested 16 RNA triggers each, since they represented 99.6% of all DENV genomes. Sensor 6 shows a good response to 15 out of 16 tested triggers. whereas sensor 7 responds to 14 out of 16 tested triggers. Trigger DENV 130 that is not sensed by sensor 7 accounts for only 5 genomes and DENV134 that is not sensed by either sensor 6 or 7 is present only in 1 genome, hence do not represent a significant genome population. Together, these results confirm that the toehold sensors do respond to even 3 mismatches in the trigger. Importantly, sensors 3, 6 and 7 are each able to detect nearly 99.4% all DENV genomes including all known serotypes (Fig S5 A-C). Hence these are truly *pan-dengue* sensors.

**Figure 4:**
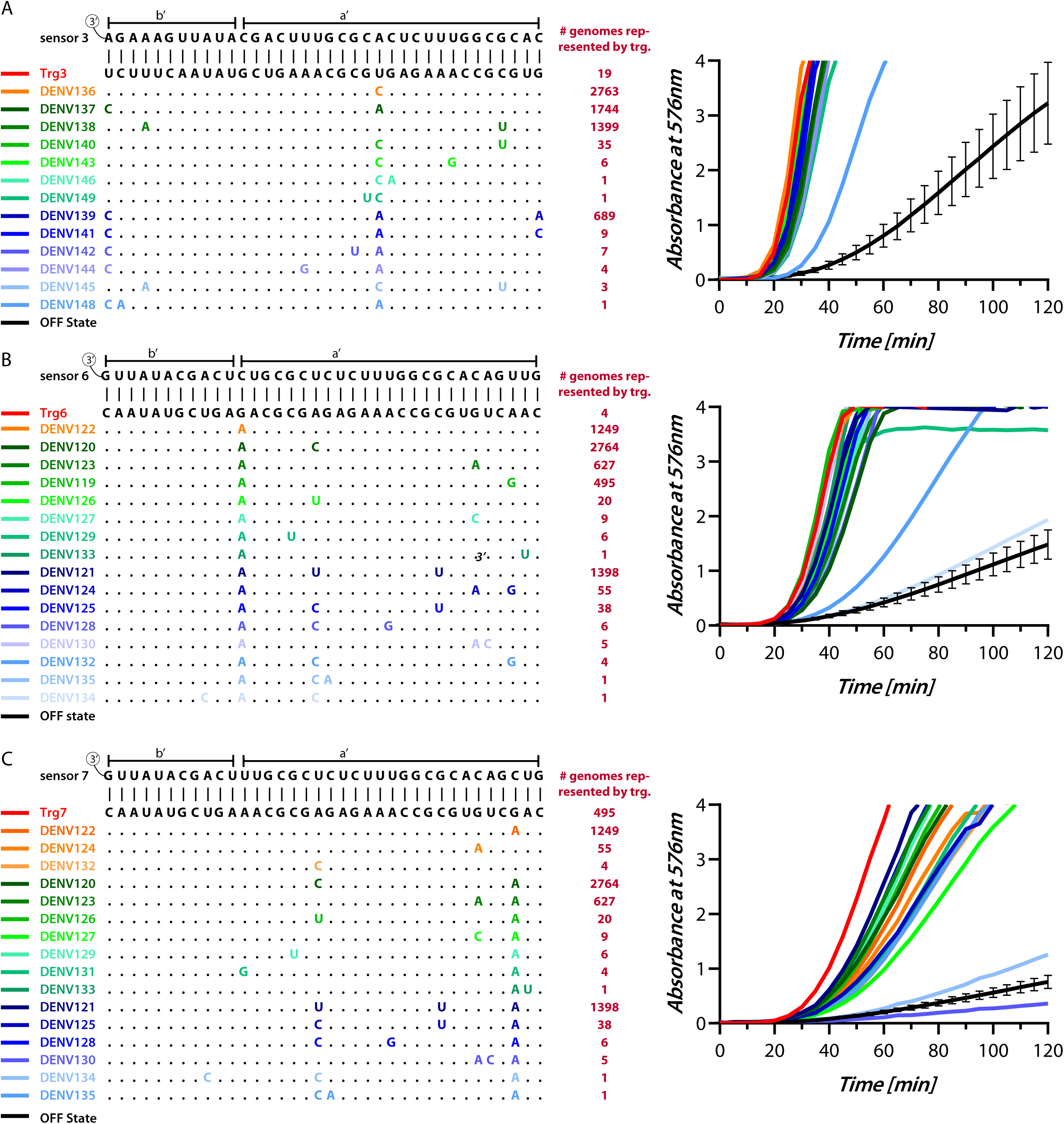
Detection of different DENV serotypes by toehold sensors. Left panel shows Multiple sequence alignment of different triggers in a trigger pool with 0 to 3 mismatches (highlighted in color) from the parent trigger (black). Also shown are the trigger binding regions of the respective sensors 3 (panel A), 6 (B), and 7 (C). The number of genomes that contain each trigger sequence is shown (red). Right panel shows the IVTT assay performed with a sensor in the absence (black curve is an average of all off states, with error bars representing standard deviation) or presence of various triggers from the trigger pool (colored lines, colors are matched with the left panel). IVTT was monitored through change in absorbance at 576 nm with 10^13^ copies of each trigger. All three sensors detect the majority of triggers tested, accounting for >99.4% of all DENV genomes. These results show that sensors 3, 6, and 7, are *pan-dengue*.

### Coupling of sensors with isothermal RNA amplification

A reported feature of all toehold sensors, including our *pan-dengue* sensors (Fig S6) is that their limit of detection is in the order of 10^12^ to 10^13^ copies of trigger RNA^32–34,36,38,40^. To increase sensitivity, toehold sensor assays are typically coupled with a step of target RNA amplification like RT-LAMP^34^ or NASBA^32,33,36,38^. We noted that sequence diversity in viral genomes may affect the amplification step that precedes toehold-based detection, and hence attempted to couple our *pan-dengue* sensor with an isothermal amplification strategy that would account for viral sequence diversity.

To this end, we attempted to design NASBA primers that could amplify the regions around and including the 36-nucleotide trigger. As expected, we were unable to find any one pair of NASBA primers that could take into account the significant sequence variations within DENV genomes. Hence, we resorted to multiple primer sets that would amplify the trigger region of most genomes of a serotype, but not limited to it. To test this we first synthesized short template RNAs representative of DENV-1, 2, 3, and 4. These RNAs were used as templates in NASBA reactions with respective primer pairs (see Methods). NASBA is an isothermal amplification method where the reverse primer initiates the synthesis of the cDNA strand. RNAse H degrades the RNA from the RNA:DNA hybrid, and this is followed by second strand synthesis by a forward primer containing the T7 promoter sequence. T7 polymerase in the reaction, transcribes RNA corresponding to the region between the forward and reverse primers. This RNA serves as a template for iterative amplification of the target RNA^46^. In our experiment, NASBA was initiated with 10^8^ copies of template RNA. Upon completion, NASBA reactions were tested for their ability to turn on sensor 3 in IVTT assays. We found that sensor 3 shows a reproducible increase in absorbance in the presence of the NASBA reaction as compared to a “no-template” control where NASBA was performed without any template RNA (Fig 5, Fig S7). Notably, 10^8^ copies of trigger RNA by themselves were unable to elicit a response from sensor 3, but post-NASBA the target RNA was amplified to a detectable range. This indicates that our *pan-dengue* sensor 3 is efficiently coupled to NASBA and that the primer sets that we have designed are efficient in amplifying targets from all serotypes.

**Figure 5:**
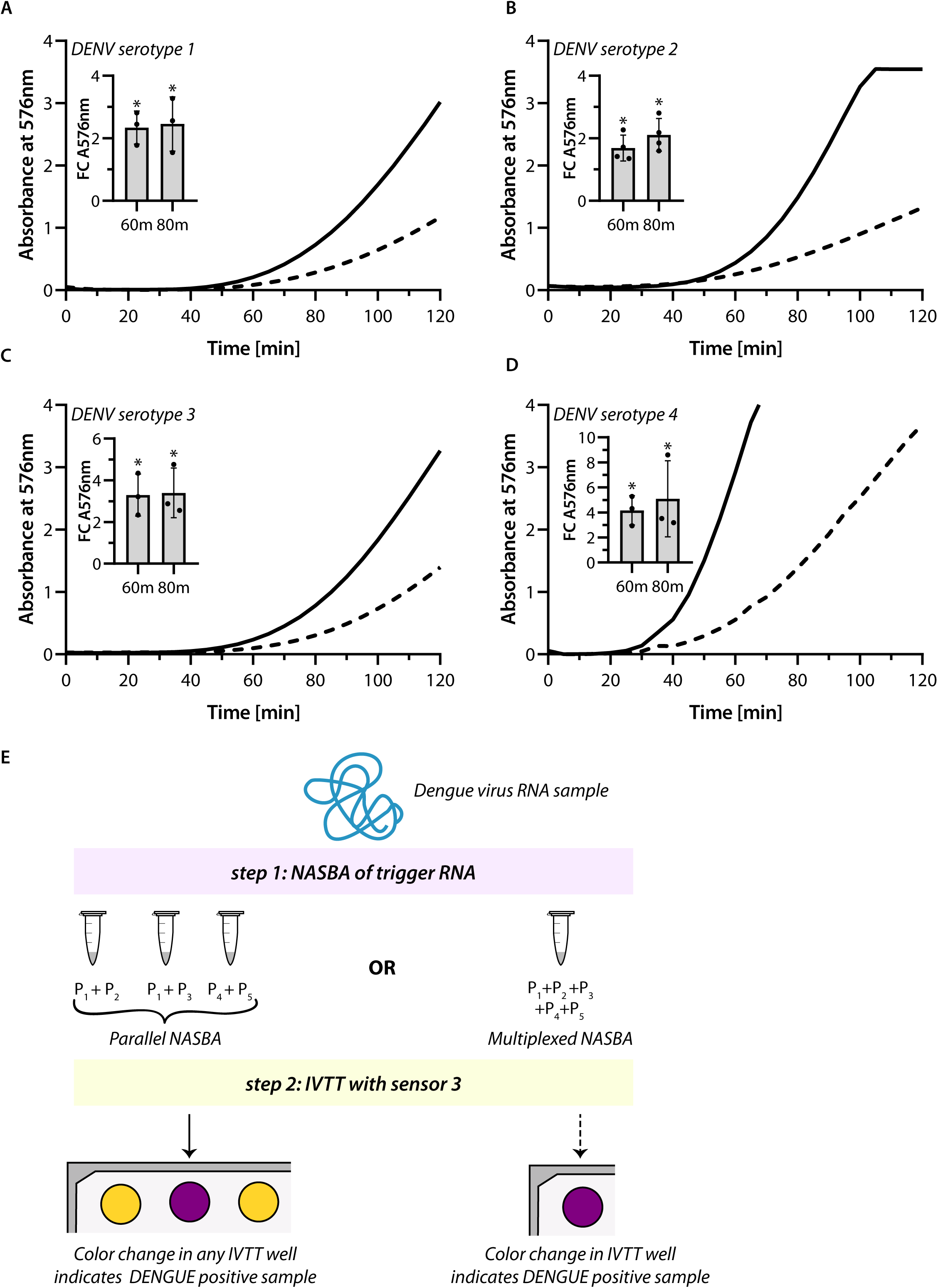
Coupling of sensors with isothermal RNA amplification. NASBA reactions were performed on templates representing different serotypes-DENV Serotype 1 (A), DENV Serotype 2 (B), DENV Serotype 3 (C), and DENV Serotype 4 (D). NASBA reactions post 2 hours were added to an IVTT reaction containing sensor 3. Change in absorbance at 576 nm was recorded over time. NASBA reactions performed on the different templates (solid lines) were compared with reactions where no template was added to the NASBA (dashed lines). Fold change in absorbance in the NASBA reactions at 60 and 80 minutes compared to the NTC is shown in inset. Inset: Bars represent mean and error bars represent standard deviation in the fold change in three or more replicates. Statistical significance was determined through unpaired t-tests. (* p < 0.05) E) Fold change in absorbance at 576 nm with respect to the NTC for each NASBA reaction. Bars indicate mean fold change, error bars indicate standard deviation, and circles indicate the fold change in each replicate. Data from three or more replicates is shown. F) Schematic shows how our *pan-dengue* sensors may be combined with different NASBA primer sets (P1, P2, P3, P4, P5) to enable DENV diagnostics. In one scheme, individual NASBA reactions each containing one primer pair would be performed and followed by IVTT with sensor 3. Assay that changes color from yellow to purple would indicate presence of DENV virus in the sample. An alternate scheme would be where a single NASBA reaction is performed where all 5 primers are multiplexed and this is followed by IVTT to detect presence (purple) of virus in the sample.

Based on these results we propose that facile detection of DENV, a virus that shows very high sequence divergence, would be possible using toehold switches. To make this possible, NASBA using multiple primer pairs would be coupled with a *pan-dengue* sensor in individual or potentially multiplexed reactions.

## DISCUSSION

In this work, we report a strategy to develop a toehold sensor assay, for a highly evolved virus such as DENV. Our analysis of the DENV genome revealed significantly high sequence divergence even among the different serotypes of DENV. Since this posed a challenge in developing *pan-dengue* sensors, we came up with a strategy where we first take into account the sequence diversity of all available genomes and select targetable regions common to them. This alone was not enough and we added another step of allowing 0 to 3 mismatches between the target and sensor. This allowance led us to identify the most conserved and hence targetable region within DENV genomes. Based on this we designed toehold sensors, which when filtered through established toehold criteria yielded three *pan-dengue* sensors that we show can detect close to 99.4% of all known dengue genomes including all serotypes. We found these sensors could be coupled with NASBA, an isothermal amplification method. This results in more sensitive detection of all the DENV serotypes. Hence our sensors are truly *pan-dengue* and we establish a method that enables design of toehold switches for detection of highly RNA divergent viruses.

The toehold RNA switch presents a powerful method to detect viral nucleic acids since it utilizes the intermolecular interactions between the sensor and trigger RNA, resulting in the control of translation of any reporter gene that enables color or fluorescence based read-out. Designing toehold switches especially against RNA viruses can be challenging, since RNA viruses may exhibit very high levels of sequence divergence over time. This is ascribed to their replication mechanisms that typically lack proof-reading. DENV, NORO, HIV, and Lassa are examples of such viruses, which have evolved over time to accumulate a substantial degree of sequence variations^47–50^. While developing nucleic acid tests for such viruses, it would be vital that the design accounts for sequence divergence by taking into account all the available genome sequences. A recent report describes a toehold based sensor that responds to DENV RNA^51^. This work presented a strategy for increasing toehold sensitivity, however the reported sensor was designed only to detect a specific DENV genome, not addressing the diversity of prevalent DENV serotypes.

Crucial to our design strategy was that we allowed 0 to 3 mismatches between the trigger RNA and the sensor. Even with 3 mismatches we saw a clear response from all three selected sensors. This showed tremendous tolerance in the toehold’s interactions with its trigger. Triggers that were not sensed (DENV 130 by sensor 7 and DENV 134 by both sensor 6 and 7) both had 3 mismatches with their cognate sensor. DENV 134 shows low single-strandedness, coupled with an unusually high Minimal Free Energy Secondary Structure (-15.5 kcal/mol) compared to all other triggers hence it is possible that it forms stable intramolecular interactions within itself. DENV 130 bears 3 mismatches in close proximity to each other, offering a plausible reason why this is not sensed by sensor 7. A previous study designed toehold switches to distinguish between wild-type target RNA versus different Single Nucleotide Polymorphisms^52^. Here mismatches govern the opening of the sensors, further highlighting the tunability of toehold switches towards mismatches.

Our design of toehold sensors requires us to find short 36-nucleotide regions with high conservation in the genome. With this narrow criterion, we found that the most conserved region of the genome lies in the coding region of the capsid protein that along with the membrane protein and envelope protein constitute the structural proteins of DENV. This region has been shown to be involved in long range interactions with the 3’ untranslated region of the DENV genome to facilitate replication. It is possible that the importance of these interactions contributes to the high degree of conservation we see.

The motivation behind our study was to develop *pan-dengue* sensors that can detect almost all the sequence variations in DENV. While we achieved this with the sensors alone, we needed to couple our sensors with isothermal amplification (NASBA) for sensitive detection. This posed a problem in terms of finding conserved primers. Hence, we had to select primers from different serotypes (but not exclusive to each serotype). These primers showed sensitive detection across all 4 DENV serotypes and worked well when coupled with sensor 3. We envision a diagnostic where these primers may be multiplexed in a single tube during amplification or can be used as individual primer sets in parallel reaction, followed by detection via our *pan-dengue* sensor (Fig 5E).

## METHODS

### Estimating nucleotide level diversity in RNA viruses

Sequences of DENV (taxon lineage ID: 12637), ZIKV (taxon lineage ID: 64320), MERS (taxon lineage ID: 1335626), and NORO (taxon lineage ID: 142786), which were marked as complete sequences and isolated from human host, were downloaded from the BV-BRC Database (Pickett et al., 2012) in FASTA (Lipman and Pearson, 1985, Pearson and Lipman, 1988) format, along with the corresponding metadata. The genomes were indexed and aligned using the Augur toolkit (Hadfield et al., 2018) with NC_035889.1, NC_038294.1, NC_044853.1, and NC_001474.2 as the reference genomes for Zika, MERS, Noro, and Dengue viruses respectively.

The alignments were analyzed through an in-house code where only positions with occupancy greater than 50% were considered for the analysis. Shannon entropy for a given position x, was calculated using the formula: *Η_x_* =-_∑_ *p_n, x_* ln *p_n, x_* where *p_n, x_* is the fraction of genomes that have a particular nucleotide *n* at position *x*.

### Analysis of Dengue genomes and identification of conserved regions in Dengue genomes

Genomes of DENV which were qualified as full-length and isolated from human sources were downloaded from the BV-BRC Database (Pickett et al., 2012) in FASTA (Lipman and Pearson, 1985, Pearson and Lipman, 1988) format on October 20, 2021. 6,716 in total. Genomes with more than 5% low-quality positions were discarded to obtain 6,712 genomes. The genomes were grouped into their respective serotypes Dengue 1 (2595 genomes), Dengue 2 (1714 genomes), Dengue 3 (1281 genomes), and Dengue 4 (543 genomes) according to the database; sequences with no serotype assignment, were assigned to Unclassified- “UC” (579 genomes). The multi-line genome sequence entries were appended to give a single-line genome sequence for every entry. All the genome entries along with their respective headers from each serotype were concatenated and placed under a line that started with “#” followed by the serotype specifier label.

Using a custom script, we identified all possible thirty-six nucleotide segments from all genomes in every serotype. Sequences that were composed of only the standard four nucleotides were considered. The occurrences of each trigger in all the DENV genomes was calculated. We applied a *Similarity Criteria* wherein we took triggers which showed 0 to 3 mismatches as compared to a parent trigger and pooled them together (see representative triggers in Fig Table 1). Each tigger in a trigger pool was analyzed for its occurrence across genomes of every serotype. This gave an estimate of the prevalence of each trigger pool across each serotype. We selected only 61 trigger pools, since they were prevalent in >95% of genomes of every serotype.

### Design of Toehold RNA sensors

Toehold sensors were designed as described in *Chakravarthy et al., 2021*. 61 trigger pools were taken, and the parent trigger of each pool was used to design a corresponding toehold sensor. The design includes a single nucleotide position between the toehold-lock (b) region and the linker (see Fig 1C). This single nucleotide position is an A, U. G or C, resulting in four possible sensors for each parent trigger. Any toehold sensors that have a stop codon are discarded. This resulted in 219 sensors in total.

We then calculated trigger single strandedness, toehold single strandedness, probability of formation of the lower stem pairing and similarity to expected sensor secondary structure. For trigger single strandedness, toehold single strandedness calculations we used the *complexdefect* module of Nupack 3.2.2 (Zadeh et al., 2011). For probability of formation of the lower stem pairing and similarity to expected sensor secondary structure we used the *pairs* module of Nupack 3.2.2. From the 219 sensors designed above, we obtained 67 sensors that met arbitrary criteria of Trigger SS>0.49, Toehold SS>0.32, lower stem score >0.97 and expected secondary structure score >0.74.

The specificity of the chosen triggers was assessed through nucleotide BLAST with the *blastn* algorithm against the non-redundant database excluding Dengue virus (taxid: 12637). Triggers were analyzed for their similarities (e < 1) to human or other related pathogens.

### *In vitro* transcription coupled translation assay (IVTT)

The IVTT reaction conditions were adapted from Chakravarthy et al., 2021^33^. The assay was conducted using the NEB PURExpress kit (Cat. no. E6800L). The reaction mixture consisted of 2 μl of solution A, 1.5 μl of solution B, 0.125 μl (10 Units) of RNase Inhibitor (Thermo Fisher Scientific, Cat. no. 10777019), ChlorophenolRed-β-D-galactopyranoside-CPRG (Sigma-Aldrich, Cat. no. 59767) 0.375 μl of 12 mg/ml dye was added as a substrate, and 125 ng linear DNA template of the sensor fused to *lacZ* gene. The IVTT reactions were incubated at 37°C in 384-well plates (Corning, Cat. no. 3544) with 1.25 μl of the cognate 9 μM Trigger RNA or, 1.25 μl of NASBA products or, indicated copies of trigger RNA or, 1.25 μl of nuclease free water. The absorbance at 576 nm was monitored every 5 minutes for 2 hours in a Varioskan Lux instrument (Thermo Fisher Scientific) or Tecan Infinite M Plex, or the reactions were quenched at the end of 2 hours with 2 μl of 2 M Na_2_CO_3_ and the absorbance of the ten-fold diluted samples were recorded using the Eppendorf Biospectrometer with an Eppendorf µCuvette G1.0. The fold change in absorbance was calculated relative to the reaction where only water was added, sensor “OFF” state. All absorbance-based plate reader experiments were initially baseline corrected using a blank sample, and each sample was normalized so that the lowest absorbance measurement was set to 0. Data visualization and figure creation were carried out using GraphPad Prism 8 and Adobe Illustrator, respectively.

### Preparation of trigger and template RNA

Trigger and template RNA for the IVTT and NASBA were prepared through in vitro transcription of corresponding DNA templates that had a T7 promoter. DNA templates of the triggers were prepared by amplifying the template oligomers with the corresponding forward and reverse primer specified in Table S2 for triggers used in Fig. 3B or Table S3 for triggers used in Fig. 4A-C. The template RNA corresponding to DENV serotype 1 and 3 were constructed through a PCR that involved amplification of two long oligomers that were partially complementary, by corresponding forward and reverse primers specified in Table S4.

Template RNA corresponding to DENV serotype 2 and 4 were constructed in two steps, and the final sequence was encoded by two disjoint long oligomers. The first steps involved extending the long oligomers to incorporate elements that are complementary to the other long oligomers using primers mentioned in Table S4. Finally, the purified products of the previous extension PCR were added in stoichiometrically equal amounts and amplified by primers specified in Table S4.

The RNA templates for NASBA reactions and Trigger RNAs for cell free IVTT reactions were synthesized by in vitro transcription reactions. This was done in a 40 μl in vitro transcription reaction system. Each reaction contained 1 μg of the relevant DNA template, 4 μl of 10X T7 polymerase reaction buffer (Toyobo, Cat. no. TRL-201), 5.5 μl of 50 mM MgCl_2_, 4 μl of 25 mM rNTPs (NEB, Cat. no. N0450S), 2 μl (100 units) of T7 RNA polymerase enzyme (Toyobo, Cat. no. TRL-201), 2 μl (0.2 units) Yeast Inorganic Pyrophosphatase (NEB, Cat. no. M2403S), 0.5 μl (20 units) RNaseOUT (Invitrogen, Cat. no. 10777019), and the remainder of the reaction volume was made up to 40 μl with nuclease free water and incubated at 37°C for 2 h. After this, the samples were treated with 2 μl (4 units) of DNAse I enzyme (NEB, Cat. no. M0303S) at 37°C for 1 hour and purified using the ZymoResearch RNA Clean and Concentrator RNA purification kit (Cat. no. 1015). The final RNA sample was eluted in nuclease free water for further use.

## NASBA

The NASBA reaction performed as described previously Chakravarthy et al., 2021^33^. The reactions were carried out by adding 10^8^ copies of template RNA with a master mix containing 4 μl of 5X AMV RT Buffer (Promega, Cat. No. M515A or Roche, Cat. no. 10109118001), 1.6 μl of 50 mM MgCl_2_, 2 μl of 25 mM rNTPs (NEB, Cat. no. N0450S), 2 μl of 10 mM dNTPs (NEB, Cat. no. N0447S), 0.18 μl of 1 M DTT (VWR, Cat. no. 3483-12-3), 3 μl of 100% DMSO (Sigma-Aldrich, Cat. no. D8418-50 Ml), 0.2 μl of 10 mg/ml BSA (Roche, Cat. no. 10735078001), and 0.5 μl each of 10 μM forward and reverse primers. The reaction mixture was made up to 17.7 μl, and the contents were heated at 65°C for 5 minutes, followed by incubation at 50°C for an additional 5 minutes. An enzyme mix containing 1 μl (50 Units) T7 RNA Polymerase (Toyobo, Cat. no. TRL-201), 1 μl (20 Units) AMV-RT (Promega, Cat. No. M510F or Roche, Cat. no. 10109118001), 0.1 μl (0.2 Units) RNaseH (Roche, Cat. no. 10786357001), and 0.2 μl (12.5 Units) of RNaseOUT (Invitrogen, Cat. no. 10777019) was added to the reaction, and incubated at 42°C for 2 hours. The reactions were stored at -80°C or used in IVTT assays. The RNA templates used in the reactions for DENV serotypes 1-4 are declared in Table S4. The primers together covered 90% of the DENV genomes that reported at least 100 nucleotides upstream of trigger pool 3 sequences (3964 genomes). Sequences of all primers used in this study are declared in Table S5.

## Acknowledgements

We thank Prof. Rakesh Mishra for supporting and funding this project. We acknowledge the support from National Centre for Biological Sciences-Tata Institute of Fundamental Research (TIFR), under project no. 12-R&D-TFR-5.040800 to AR. We thank Institute of Eminence Directorate, University of Hyderabad (UoH) for funding, including grant UoH-IoE-RC3-21-035 funding to AG. We thank Department of Biochemistry, School of Life Sciences (SLS) and UoH for support, and DBT, India for DBT-SAHAJ/BUILDER grant # BT/INF/22/SP41176/2020 to SLS. We are grateful to DST-FIST 2023 program grant SR/FST/LS-II/2023/1172 for funding SLS. AN is supported by a DBT-JRF/SRF fellowship (DBT/2020/NCBS/1442).

## Author Contributions

AN, DM, AS, and EM performed experiments, analyzed data, and contributed to the writing of the manuscript. AR and AG conceptualized the study, designed the methodology, supervised the work, administered the project, acquired funding, wrote the original draft, and reviewed and edited the writing.

## Data, Materials, and Software Availability

All study data are included in the article and/or supporting information. Any other information would be available upon request from the authors.

## Supplementary legends

**Figure S1:**
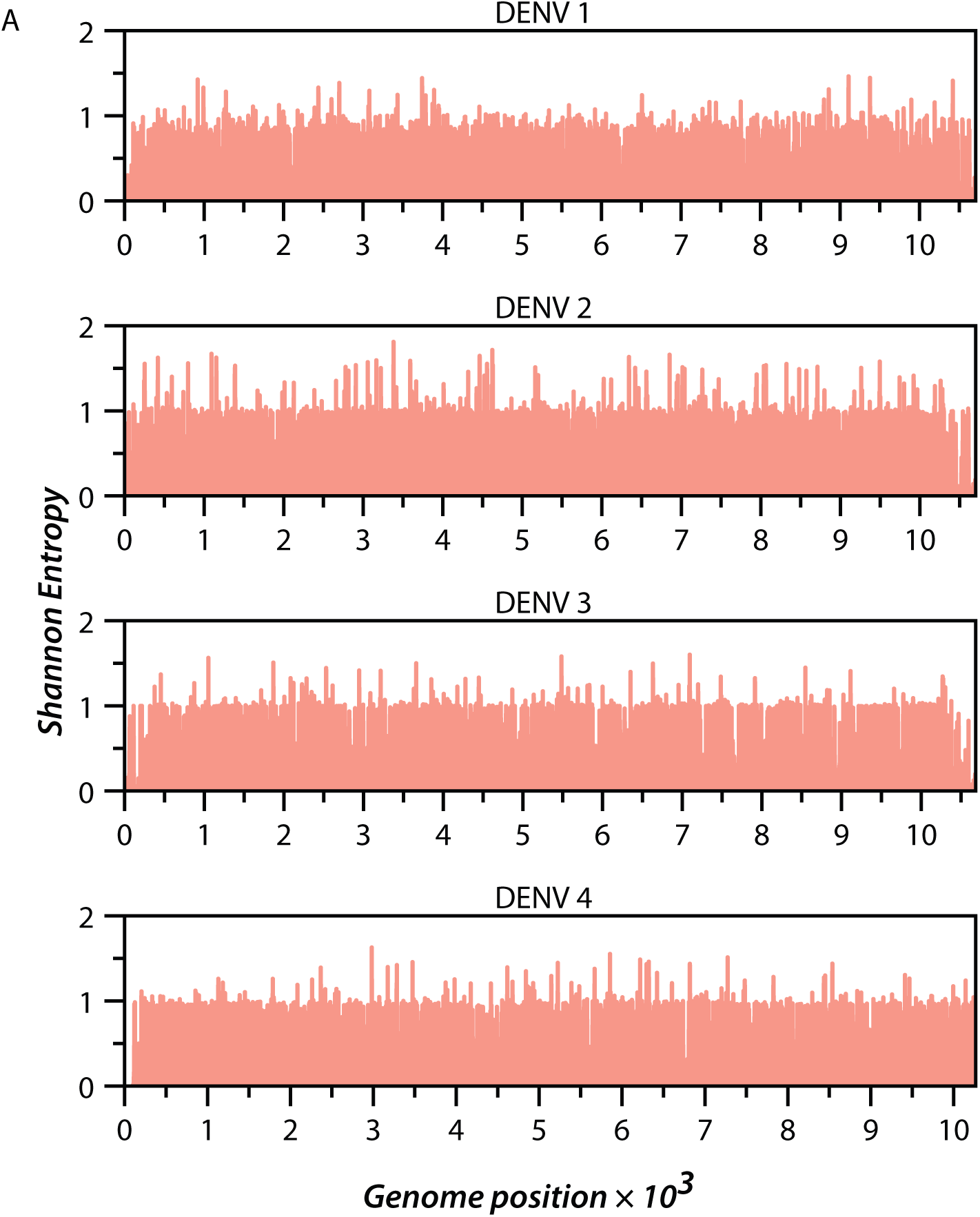
Extent of diversity within genomes of each DENV serotype. A) Diversity in the composition of nucleotides at each position across the genomes of DENV serotypes 1-4 is shown in terms of Shannon entropy.

**Figure S2:**
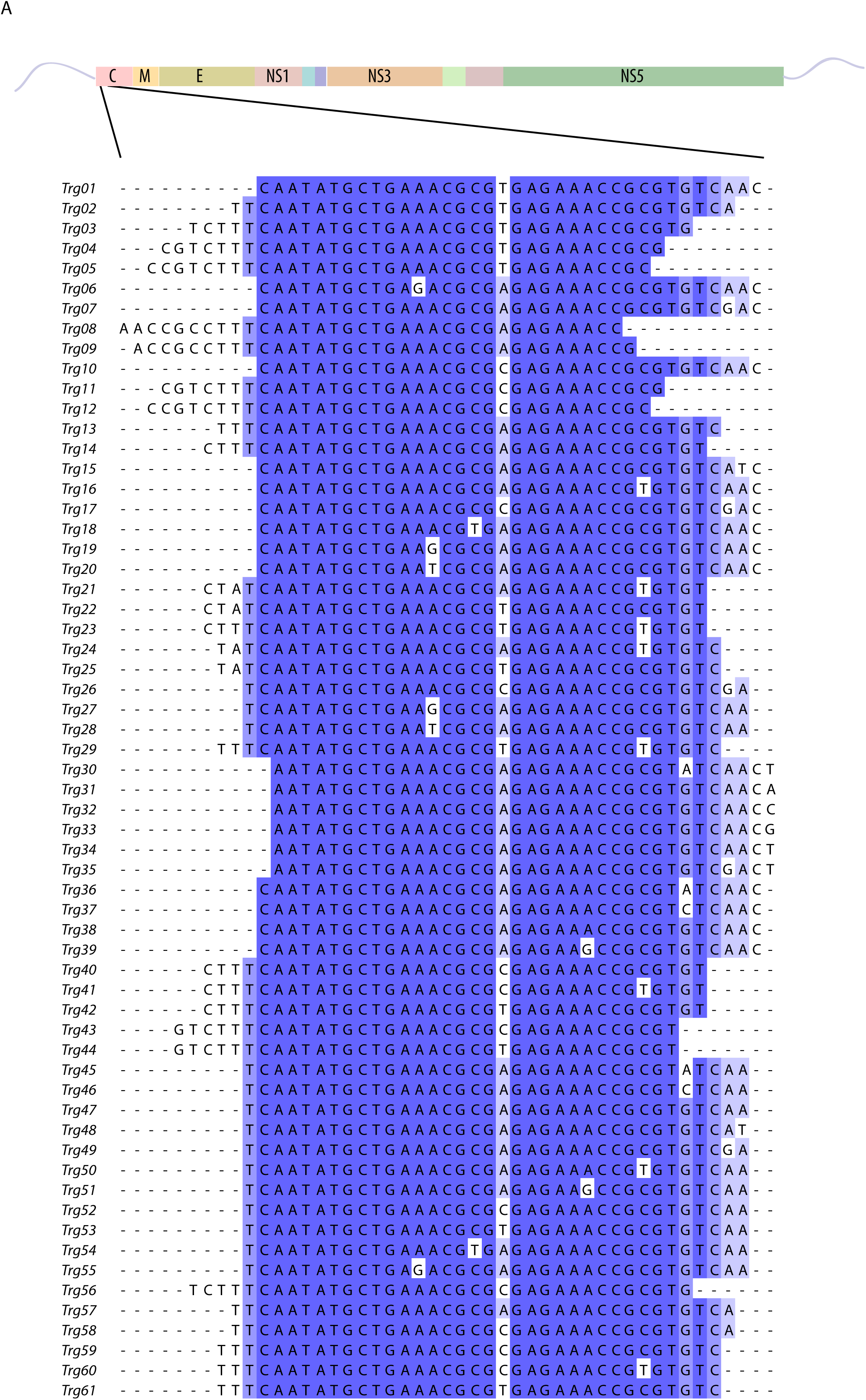
All 61 *pan-dengue* trigger pools identified in this study are located in the Capsid protein coding region of the DENV genome (top). Sequences of the parent trigger of all 61 *pan-dengue* trigger pools were aligned using MAFFT^53^ and have been shaded on the basis of nucleotide conservation. Darker shades indicate higher nucleotide conservation.

**Figure S3:**
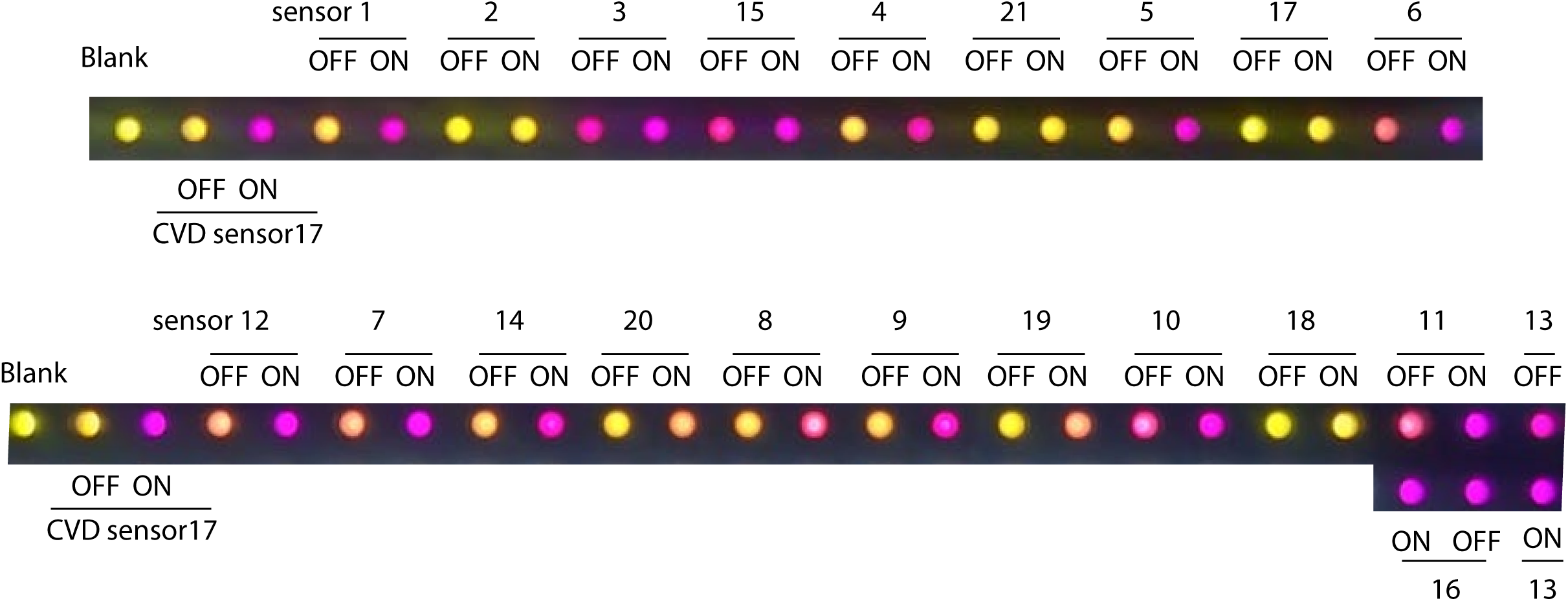
*In vitro* screening of DENV Toehold RNA sensors. In vitro transcription translation (IVTT) assay of 21 sensors in presence (ON) or absence (OFF) of their cognate triggers was performed. CPRG, a yellow-coloured substrate of LacZ added to the reaction, gets hydrolyzed by LacZ and appears purple. Cell phone camera was used to capture the image of the multi-well plate at the end of 2h at 37℃. Absorbance at 576 nm is reported in Fig 3. IVTT with no sensor or trigger (Blank) and with a previously reported SARS-COV2 toehold sensor without (CVD sensor17-OFF) and with its cognate trigger (CVD sensor 17-ON) are used as controls.

**Figure S4:**
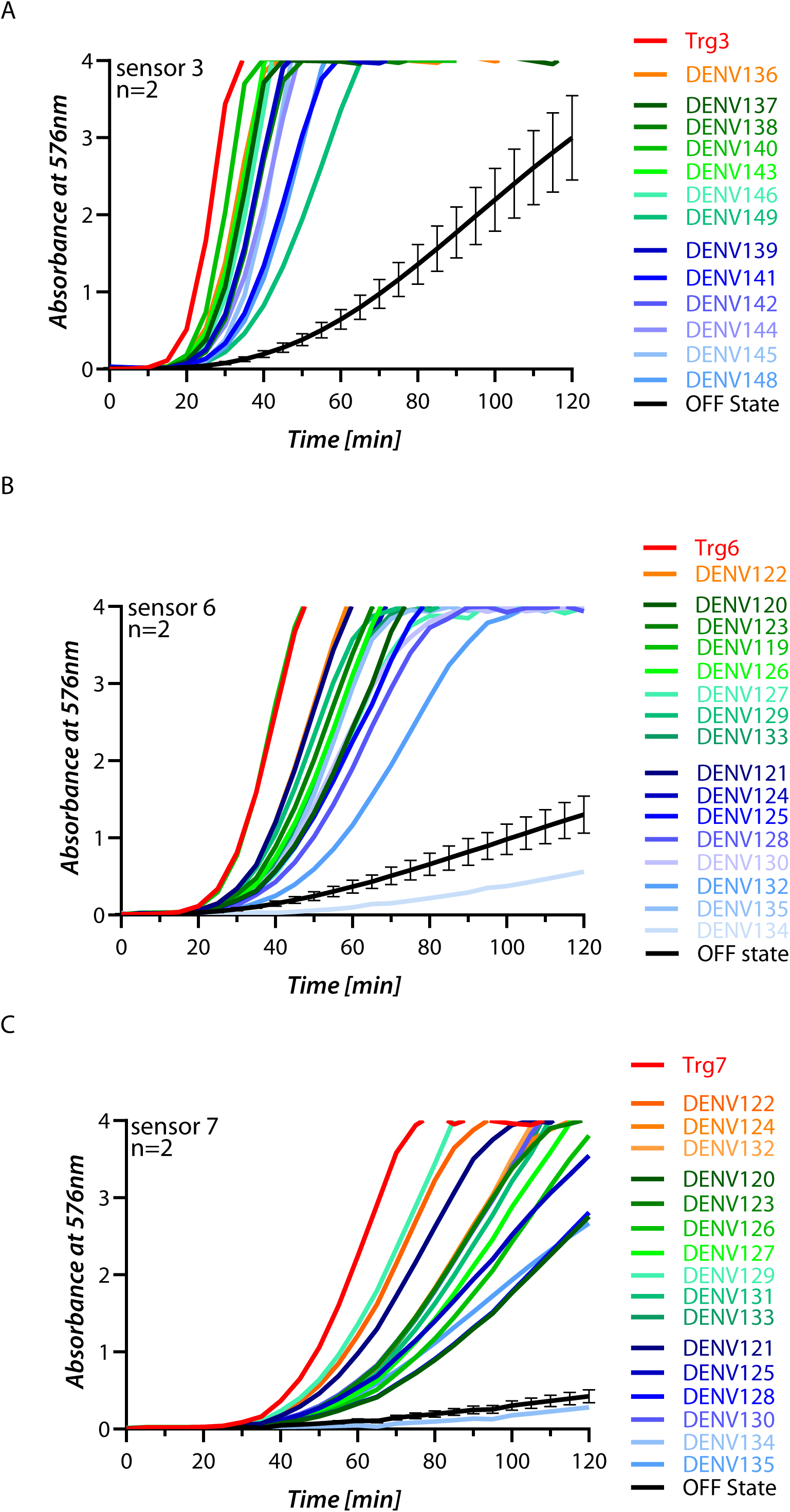
IVTT assay performed with sensors 3, 6 and 7 in the absence (black curve) or presence of various triggers from the trigger pool (colored lines, colors are matched with the left panel). IVTT was monitored through change in absorbance at 576 nm with 10^13^ copies of each trigger. N=2 replicates of the kinetics profile are shown here (N=1 shown in Figure 4 A-C).

**Figure S5:**
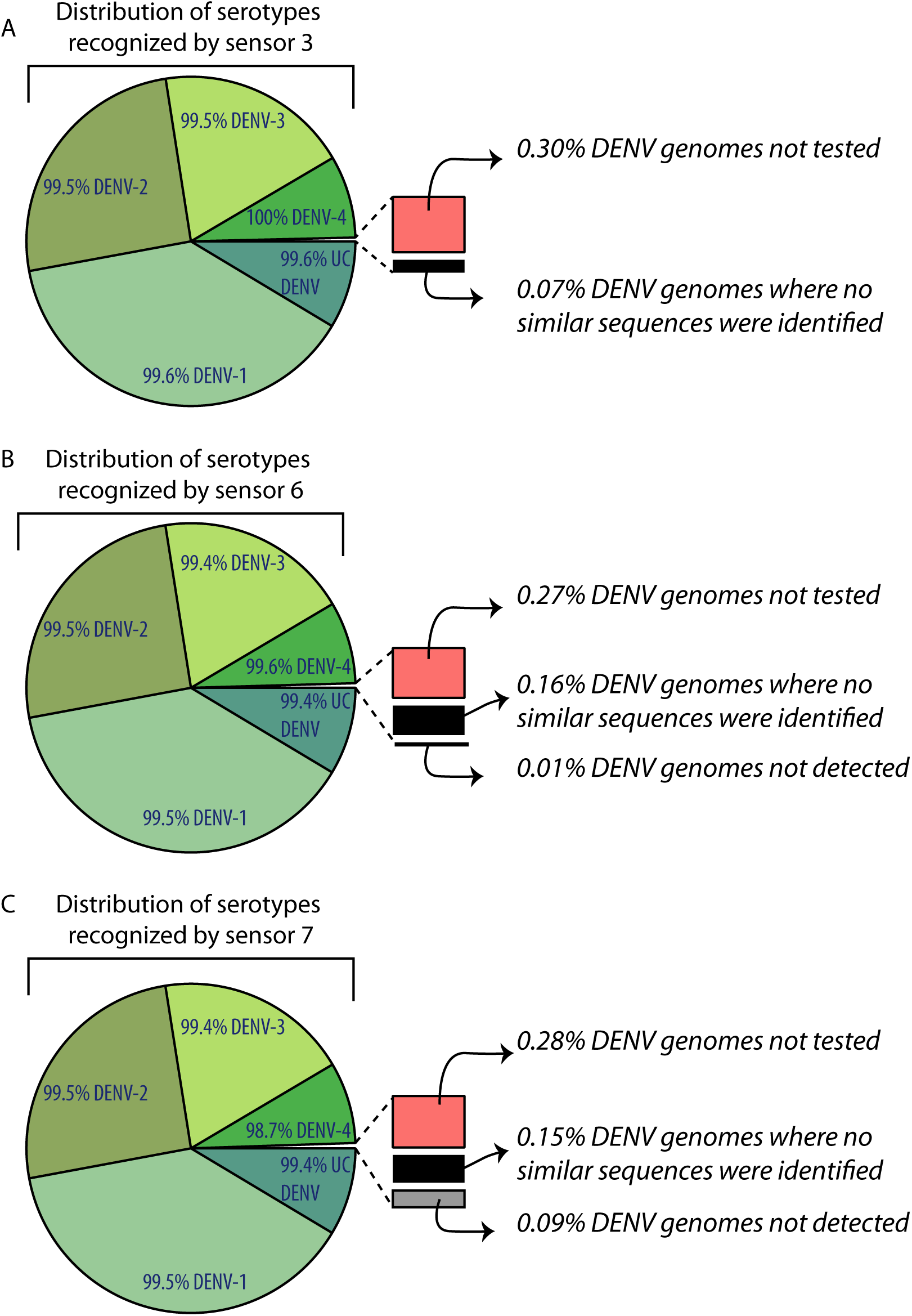
Pie-chart illustrating the distribution of DENV serotypes detected (shades of green) by our *pan-dengue* sensors 3 (A), 6 (B), and 7 (C). Slice of the pie-chart highlights genomes in which no triggers were identified (black), genomes whose Triggers were not tested (red), and genomes that were tested but not detected by the sensor (gray)

**Figure S6:**
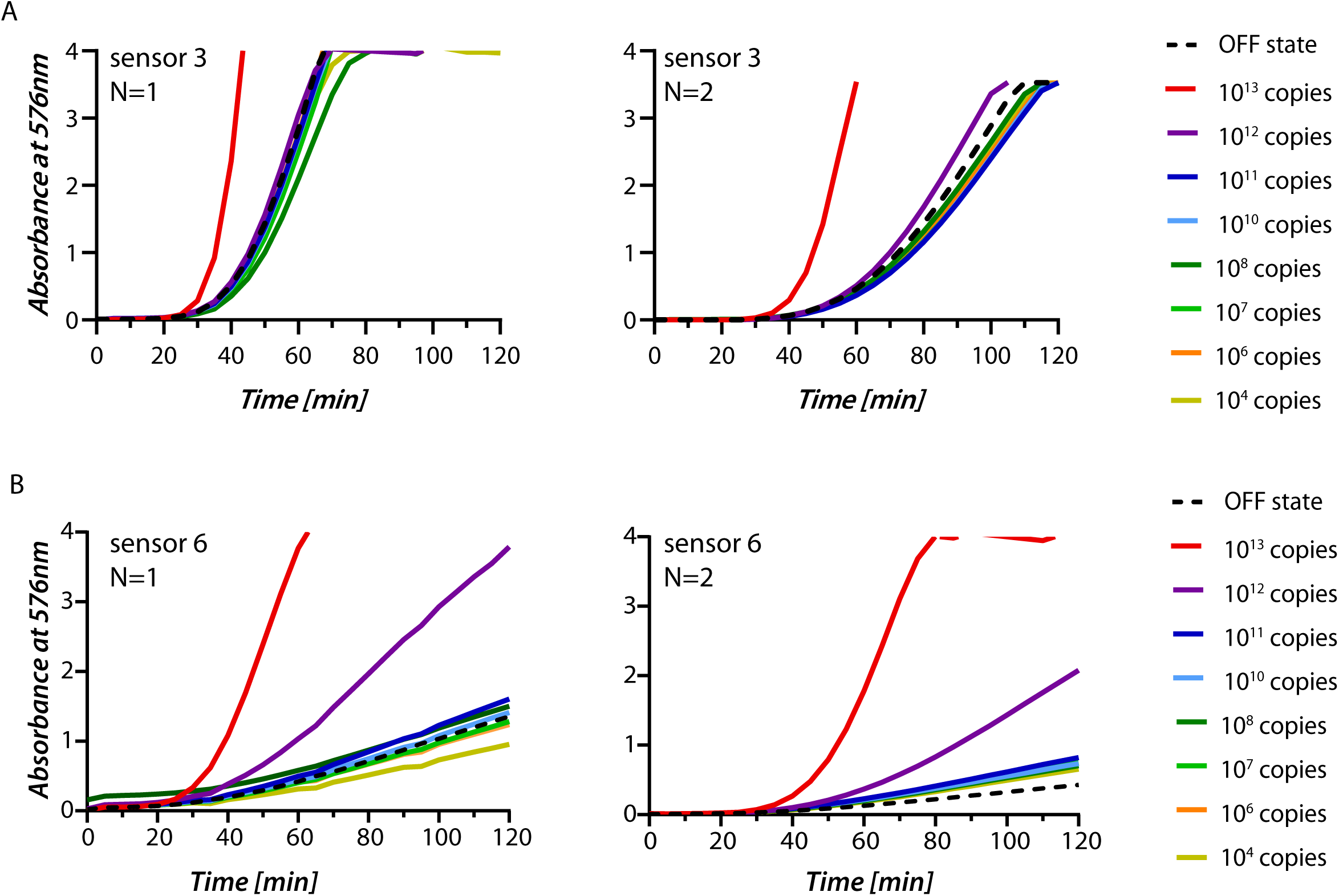
Change in absorbance over time observed in IVTT assays performed with sensor 3 (A) and 6 (B) with varying amounts of their cognate trigger RNA (ranging from 10^13^ copies to 10^4^ copies of RNA). Off state with no trigger is shown as a black dashed line. N=2 replicates are shown.

**Figure S7:**
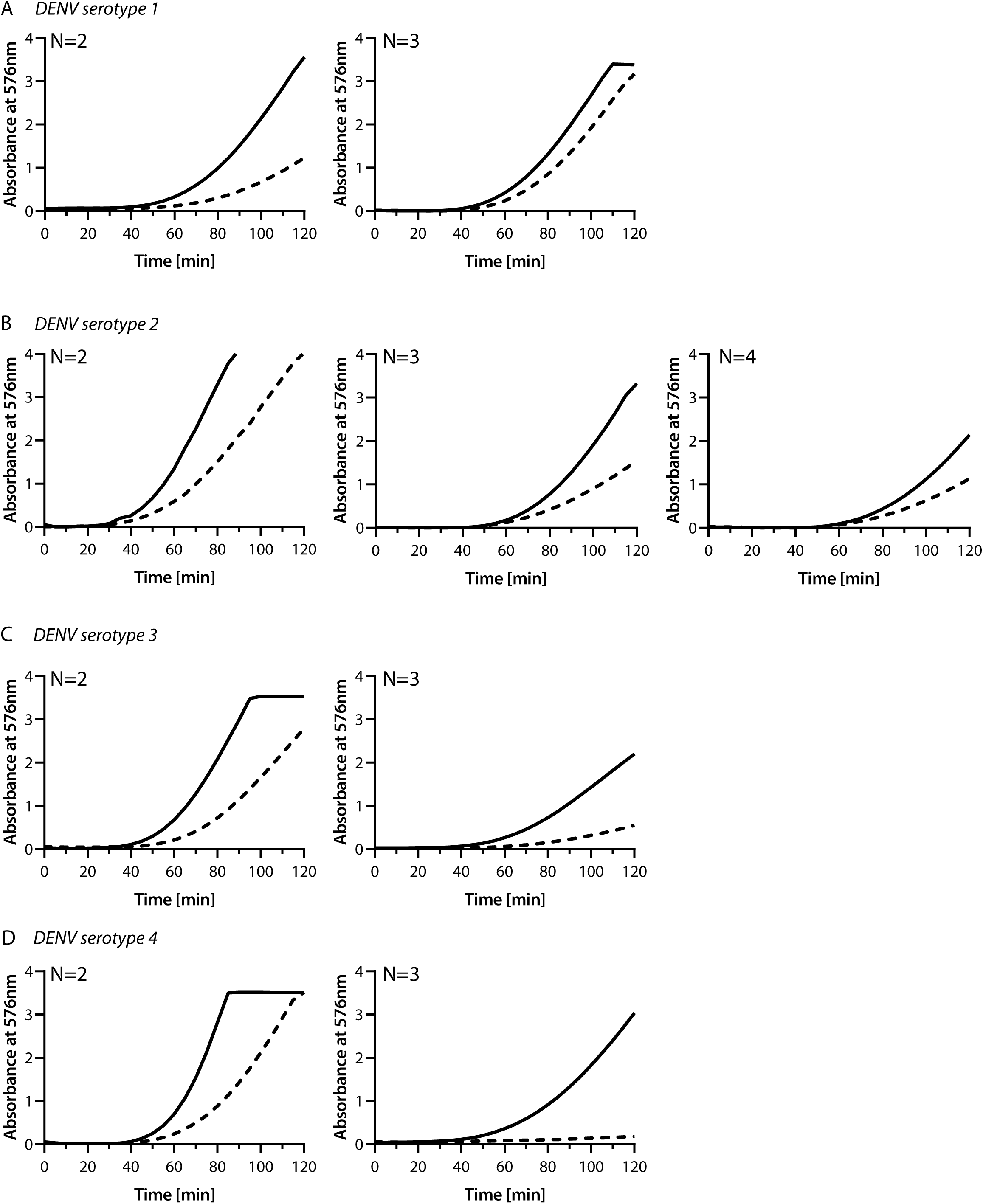
NASBA reactions were performed on templates representing different serotypes-DENV Serotype 1 (A), DENV Serotype 2 (B), DENV Serotype 3 (C), and DENV Serotype 4 (D). NASBA reactions post 2 hours were added to an IVTT reaction containing sensor 3. Change in absorbance at 576 nm was recorded over time. NASBA reactions performed on the different templates (solid lines) were compared with reactions where no template was added to the NASBA (dashed lines). N=2, 3 (or 4 where applicable). N=1 is shown in Fig. 5 A-D.

## Notes

### Competing Interest Statement

The authors have declared no competing interest.

## References

(1) Drake, J. W. Rates of Spontaneous Mutation among RNA Viruses. Proceedings of the National Academy of Sciences 1993, 90 (9), 4171–4175. 10.1073/PNAS.90.9.4171.

(2) Worobey, M.; Rambaut, A.; Holmes, E. C. Widespread Intra-Serotype Recombination in Natural Populations of Dengue Virus. Proceedings of the National Academy of Sciences 1999, 96 (13), 7352–7357. 10.1073/pnas.96.13.7352.

(3) Schoefbaenker, M.; Günther, T.; Lorentzen, E. U.; Romberg, M.-L.; Hennies, M. T.; Neddermeyer, R.; Müller, M. M.; Mellmann, A.; Bojarzyn, C. R.; Lenz, G.; Stelljes, M.; Hrincius, E. R.; Vollenberg, R.; Ludwig, S.; Tepasse, P.-R.; Kühn, J. E. Characterisation of the Antibody-Mediated Selective Pressure Driving Intra-Host Evolution of SARS-CoV-2 in Prolonged Infection. PLoS Pathog 2024, 20 (10), e1012624. 10.1371/journal.ppat.1012624.

(4) Domingo, E.; Sheldon, J.; Perales, C. Viral Quasispecies Evolution. Microbiol Mol Biol Rev 2012, 76 (2), 159–216. 10.1128/MMBR.05023-11.

(5) Sanjuán, R.; Nebot, M. R.; Chirico, N.; Mansky, L. M.; Belshaw, R. Viral Mutation Rates. J Virol 2010, 84 (19), 9733–9748. 10.1128/JVI.00694-10/ASSET/884A4C8C-706C-4F69-9F3A-450FB3FAC0BD/ASSETS/GRAPHIC/ZJV9990937050002.JPEG.

(6) Peck, K. M.; Lauring, A. S. Complexities of Viral Mutation Rates. J Virol 2018, 92 (14). 10.1128/JVI.01031-17.

(7) Hammon, W. M. D.; Rudnick, A.; Sather, G. E. Viruses Associated with Epidemic Hemorrhagic Fevers of the Philippines and Thailand. Science (1979) 1960, 131 (3407), 1102–1103. 10.1126/SCIENCE.131.3407.1102.

(8) Hotta, S. Experimental Studies on Dengue: I. Isolation, Identification and Modification of the Virus. J Infect Dis 1952, 90 (1), 1–9. 10.1093/INFDIS/90.1.1.

(9) Sabin, A. B. Research on Dengue during World War II. Am J Trop Med Hyg 1952, 1 (1), 30–50. 10.4269/AJTMH.1952.1.30.

(10) Dengue and severe dengue. https://www.who.int/news-room/fact-sheets/detail/dengue-and-severe-dengue (accessed 2025-05-24).

(11) Brady, O. J.; Gething, P. W.; Bhatt, S.; Messina, J. P.; Brownstein, J. S.; Hoen, A. G.; Moyes, C. L.; Farlow, A. W.; Scott, T. W.; Hay, S. I. Refining the Global Spatial Limits of Dengue Virus Transmission by Evidence-Based Consensus. PLoS Negl Trop Dis 2012, 6 (8). 10.1371/JOURNAL.PNTD.0001760.

(12) Young, P. R.; Hilditch, P. A.; Bletchly, C.; Halloran, W. An Antigen Capture Enzyme-Linked Immunosorbent Assay Reveals High Levels of the Dengue Virus Protein NS1 in the Sera of Infected Patients. J Clin Microbiol 2000, 38 (3), 1053–1057. 10.1128/JCM.38.3.1053-1057.2000/ASSET/52AE1765-FBAD-4754-9EE6-7F7D9A5D1A8D/ASSETS/GRAPHIC/JM0301090003.JPEG.

(13) Alcon, S.; Talarmin, A.; Debruyne, M.; Falconar, A.; Deubel, V.; Flamand, M. Enzyme-Linked Immunosorbent Assay Specific to Dengue Virus Type 1 Nonstructural Protein NS1 Reveals Circulation of the Antigen in the Blood during the Acute Phase of Disease in Patients Experiencing Primary or Secondary Infections. J Clin Microbiol 2002, 40 (2), 376–381. 10.1128/JCM.40.02.376-381.2002/ASSET/F88DABA0-4BDD-4530-8293-31B89A124B00/ASSETS/GRAPHIC/JM0220449003.JPEG.

(14) Ding, X.; Hu, D.; Chen, Y.; Di, B.; Jin, G.; Pan, Y.; Qiu, L.; Wang, Y.; Wen, K.; Wang, M.; Che, X. Full Serotype- and Group-Specific NS1 Capture Enzyme-Linked Immunosorbent Assay for Rapid Differential Diagnosis of Dengue Virus Infection. Clinical and Vaccine Immunology 2011, 18 (3), 430–434. 10.1128/CVI.00462-10/SUPPL_FILE/SUPPLEMENTA_TABLES_FOR_DENV_NS1_ASSAY.ZIP.

(15) Clinical Testing Guidance for Dengue | Dengue | CDC. https://www.cdc.gov/dengue/hcp/diagnosis-testing/index.html (accessed 2025-05-24).

(16) Wu, S. J. L.; Eun Mi Lee; Putvatana, R.; Shurtliff, R. N.; Porter, K. R.; Suharyono, W.; Watts, D. M.; King, C. C.; Murphy, G. S.; Hayes, C. G.; Romano, J. W. Detection of Dengue Viral RNA Using a Nucleic Acid Sequence-Based Amplification Assay. J Clin Microbiol 2001, 39 (8), 2794–2798. 10.1128/JCM.39.8.2794-2798.2001/ASSET/CB1CB97E-D196-469D-8703-6CE07E6B6F69/ASSETS/GRAPHIC/JM0810147002.JPEG.

(17) Usawattanakul, W.; Jittmittraphap, A.; Endy, T. P.; Nisalak, A.; Tapchaisri, P.; Looareesuwan, S. Rapid Detection of Dengue Viral RNA by Nuclic Acid Sequence-Based Amplification (NASBA). Dengue Bull 2002, 26, 125–130. https://iris.who.int/handle/10665/163750.

(18) Jittmittraphap, A.; Thammapalo, S.; Ratanasetyuth, N.; Wongba, N.; Mammen, M. P.; Jampangern, W. Rapid Detection of Dengue Viral RNA in Mosquitoes by Nucleic Acid-Sequence Based Amplification (NASBA). Southeast Asian J Trop Med Public Health 2006, 37 (6), 1117–1124.

(19) Abd El Wahed, A.; Patel, P.; Faye, O.; Thaloengsok, S.; Heidenreich, D.; Matangkasombut, P.; Manopwisedjaroen, K.; Sakuntabhai, A.; Sall, A. A.; Hufert, F. T.; Weidmann, M. Recombinase Polymerase Amplification Assay for Rapid Diagnostics of Dengue Infection. PLoS One 2015, 10 (6), e0129682. 10.1371/JOURNAL.PONE.0129682.

(20) Teoh, B. T.; Sam, S. S.; Tan, K. K.; Danlami, M. B.; Shu, M. H.; Johari, J.; Hooi, P. S.; Brooks, D.; Piepenburg, O.; Nentwich, O.; Wilder-Smith, A.; Franco, L.; Tenorio, A.; Abu Bakar, S. Early Detection of Dengue Virus by Use of Reverse Transcription-Recombinase Polymerase Amplification. J Clin Microbiol 2015, 53 (3), 830–837. 10.1128/JCM.02648-14/SUPPL_FILE/ZJM999094090SO6.PDF.

(21) Valloly, P.; Roy, R. Nucleic Acid Quantification with Amplicon Yield in Recombinase Polymerase Amplification. Anal Chem 2022, 94 (40), 13897–13905. 10.1021/ACS.ANALCHEM.2C02810/ASSET/IMAGES/LARGE/AC2C02810_0006.JPEG.

(22) Myhrvold, C.; Freije, C. A.; Gootenberg, J. S.; Abudayyeh, O. O.; Metsky, H. C.; Durbin, A. F.; Kellner, M. J.; Tan, A. L.; Paul, L. M.; Parham, L. A.; Garcia, K. F.; Barnes, K. G.; Chak, B.; Mondini, A.; Nogueira, M. L.; Isern, S.; Michael, S. F.; Lorenzana, I.; Yozwiak, N. L.; MacInnis, B. L.; Bosch, I.; Gehrke, L.; Zhang, F.; Sabeti, P. C. Field-Deployable Viral Diagnostics Using CRISPR-Cas13. Science (1979) 2018, 360 (6387), 444–448. 10.1126/SCIENCE.AAS8836/SUPPL_FILE/AAS8836-MYHRVOLD-SM.PDF.

(23) Gootenberg, J. S.; Abudayyeh, O. O.; Lee, J. W.; Essletzbichler, P.; Dy, A. J.; Joung, J.; Verdine, V.; Donghia, N.; Daringer, N. M.; Freije, C. A.; Myhrvold, C.; Bhattacharyya, R. P.; Livny, J.; Regev, A.; Koonin, E. V.; Hung, D. T.; Sabeti, P. C.; Collins, J. J.; Zhang, F. Nucleic Acid Detection with CRISPR-Cas13a/C2c2. Science (1979) 2017, 356 (6336), 438–442. 10.1126/SCIENCE.AAM9321/SUPPL_FILE/PAPV2.PDF.

(24) Parida, M.; Horioke, K.; Ishida, H.; Dash, P. K.; Saxena, P.; Jana, A. M.; Islam, M. A.; Inoue, S.; Hosaka, N.; Morita, K. Rapid Detection and Differentiation of Dengue Virus Serotypes by a Real-Time Reverse Transcription-Loop-Mediated Isothermal Amplification Assay. J Clin Microbiol 2005, 43 (6), 2895–2903. 10.1128/JCM.43.6.2895-2903.2005/ASSET/C301C9B1-8BAF-4199-87BD-5605399FE5F4/ASSETS/GRAPHIC/ZJM0060553950006.JPEG.

(25) Teoh, B. T.; Sam, S. S.; Tan, K. K.; Johari, J.; Danlami, M. B.; Hooi, P. S.; Md-Esa, R.; AbuBakar, S. Detection of Dengue Viruses Using Reverse Transcription-Loop-Mediated Isothermal Amplification. BMC Infect Dis 2013, 13 (1), 1–9. 10.1186/1471-2334-13-387/FIGURES/5.

(26) Lu, X.; Li, X.; Mo, Z.; Jin, F.; Wang, B.; Zhao, H.; Shan, X.; Shi, L. Rapid Identification of Chikungunya and Dengue Virus by a Real-Time Reverse Transcription-Loop-Mediated Isothermal Amplification Method. Am J Trop Med Hyg 2012, 87 (5), 947–953. 10.4269/AJTMH.2012.11-0721.

(27) Dauner, A. L.; Mitra, I.; Gilliland, T.; Seales, S.; Pal, S.; Yang, S. C.; Guevara, C.; Chen, J. H.; Liu, Y. C.; Kochel, T. J.; Wu, S. J. L. Development of a Pan-Serotype Reverse Transcription Loop-Mediated Isothermal Amplification Assay for the Detection of Dengue Virus. Diagn Microbiol Infect Dis 2015, 83 (1), 30–36. 10.1016/J.DIAGMICROBIO.2015.05.004.

(28) Neeraja, M.; Lakshmi, V.; Lavanya, V.; Priyanka, E. N.; Parida, M. M.; Dash, P. K.; Sharma, S.; Rao, P. V. L.; Reddy, G. Rapid Detection and Differentiation of Dengue Virus Serotypes by NS1 Specific Reverse Transcription Loop-Mediated Isothermal Amplification (RT-LAMP) Assay in Patients Presenting to a Tertiary Care Hospital in Hyderabad, India. J Virol Methods 2015, 211, 22–31. 10.1016/J.JVIROMET.2014.10.005.

(29) Zhou, Y.; Wan, Z.; Yang, S.; Li, Y.; Li, M.; Wang, B.; Hu, Y.; Xia, X.; Jin, X.; Yu, N.; Zhang, C. A Mismatch-Tolerant Reverse Transcription Loop-Mediated Isothermal Amplification Method and Its Application on Simultaneous Detection of All Four Serotype of Dengue Viruses. Front Microbiol 2019, 10 (MAY), 446304. 10.3389/FMICB.2019.01056/BIBTEX.

(30) Tharanga, S.; Ünlü, E. S.; Hu, Y.; Sjaugi, M. F.; Çelik, M. A.; Hekimoğlu, H.; Miotto, O.; Öncel, M. M.; Khan, A. M. DiMA: Sequence Diversity Dynamics Analyser for Viruses. Brief Bioinform 2024, 26 (1). 10.1093/BIB/BBAE607.

(31) Metsky, H. C.; Welch, N. L.; Pillai, P. P.; Haradhvala, N. J.; Rumker, L.; Mantena, S.; Zhang, Y. B.; Yang, D. K.; Ackerman, C. M.; Weller, J.; Blainey, P. C.; Myhrvold, C.; Mitzenmacher, M.; Sabeti, P. C. Designing Sensitive Viral Diagnostics with Machine Learning. Nature Biotechnology 2022 40:7 2022, 40 (7), 1123–1131. 10.1038/s41587-022-01213-5.

(32) Pardee, K.; Green, A. A.; Takahashi, M. K.; Braff, D.; Lambert, G.; Lee, J. W.; Ferrante, T.; Ma, D.; Donghia, N.; Fan, M.; Daringer, N. M.; Bosch, I.; Dudley, D. M.; O’Connor, D. H.; Gehrke, L.; Collins, J. J. Rapid, Low-Cost Detection of Zika Virus Using Programmable Biomolecular Components. Cell 2016, 165 (5), 1255–1266. 10.1016/j.cell.2016.04.059.

(33) Chakravarthy, A.; Nandakumar, A.; George, G.; Ranganathan, S.; Umashankar, S.; Shettigar, N.; Palakodeti, D.; Gulyani, A.; Ramesh, A. Engineered RNA Biosensors Enable Ultrasensitive SARS-CoV-2 Detection in a Simple Color and Luminescence Assay. Life Sci Alliance 2021, 4 (12). 10.26508/LSA.202101213.

(34) Park, S.; Lee, J. W. Detection of Coronaviruses Using RNA Toehold Switch Sensors. International Journal of Molecular Sciences 2021, Vol. 22, Page 1772 2021, 22 (4), 1772. 10.3390/IJMS22041772.

(35) Kang, X.; Zhao, C.; Chen, S.; Zhang, X.; Xue, B.; Li, C.; Wang, S.; Yang, X.; Xia, Z.; Xu, Y.; Huang, Y.; Qiu, Z.; Li, C.; Wang, J.; Pang, J.; Shen, Z. Development of a Cell-Free Toehold Switch for Hepatitis A Virus Type I on-Site Detection. Analytical Methods 2023, 15 (43), 5813–5822. 10.1039/D3AY01408H.

(36) Ma, D.; Shen, L.; Wu, K.; Diehnelt, C. W.; Green, A. A. Low-Cost Detection of Norovirus Using Paper-Based Cell-Free Systems and Synbody-Based Viral Enrichment. Synth Biol 2018, 3 (1). 10.1093/SYNBIO/YSY018.

(37) Wang, S.; Emery, N. J.; Liu, A. P. A Novel Synthetic Toehold Switch for MicroRNA Detection in Mammalian Cells. ACS Synth Biol 2019, 8 (5), 1079–1088. 10.1021/ACSSYNBIO.8B00530/ASSET/IMAGES/LARGE/SB-2018-00530A_0007.JPEG.

(38) Takahashi, M. K.; Tan, X.; Dy, A. J.; Braff, D.; Akana, R. T.; Furuta, Y.; Donghia, N.; Ananthakrishnan, A.; Collins, J. J. A Low-Cost Paper-Based Synthetic Biology Platform for Analyzing Gut Microbiota and Host Biomarkers. Nat Commun 2018, 9 (1), 3347. 10.1038/s41467-018-05864-4.

(39) Green, A. A.; Silver, P. A.; Collins, J. J.; Yin, P. Toehold Switches: De-Novo-Designed Regulators of Gene Expression. Cell 2014, 159 (4), 925–939. 10.1016/J.CELL.2014.10.002.

(40) Amalfitano, E.; Karlikow, M.; Norouzi, M.; Jaenes, K.; Cicek, S.; Masum, F.; Sadat Mousavi, P.; Guo, Y.; Tang, L.; Sydor, A.; Ma, D.; Pearson, J. D.; Trcka, D.; Pinette, M.; Ambagala, A.; Babiuk, S.; Pickering, B.; Wrana, J.; Bremner, R.; Mazzulli, T.; Sinton, D.; Brumell, J. H.; Green, A. A.; Pardee, K. A Glucose Meter Interface for Point-of-Care Gene Circuit-Based Diagnostics. Nature Communications 2021 12:1 2021, 12 (1), 1–10. 10.1038/s41467-020-20639-6.

(41) Pickett, B. E.; Greer, D. S.; Zhang, Y.; Stewart, L.; Zhou, L.; Sun, G.; Gu, Z.; Kumar, S.; Zaremba, S.; Larsen, C. N.; Jen, W.; Klem, E. B.; Scheuermann, R. H. Virus Pathogen Database and Analysis Resource (ViPR): A Comprehensive Bioinformatics Database and Analysis Resource for the Coronavirus Research Community. Viruses 2012, 4 (11), 3209–3226. 10.3390/V4113209.

(42) Shannon, C. E. A Mathematical Theory of Communication. Bell System Technical Journal 1948, 27 (3), 379–423. 10.1002/j.1538-7305.1948.tb01338.x.

(43) Schmitt, A. O.; Herzel, H. Estimating the Entropy of DNA Sequences. J Theor Biol 1997, 188 (3), 369–377. 10.1006/JTBI.1997.0493.

(44) Hahn, C. S.; Hahn, Y. S.; Rice, C. M.; Lee, E.; Dalgarno, L.; Strauss, E. G.; Strauss, J. H. Conserved Elements in the 3’ Untranslated Region of Flavivirus RNAs and Potential Cyclization Sequences. J Mol Biol 1987, 198 (1), 33–41. 10.1016/0022-2836(87)90455-4.

(45) Alvarez, D. E.; Filomatori, C. V.; Gamarnik, A. V. Functional Analysis of Dengue Virus Cyclization Sequences Located at the 5’ and 3’UTRs. Virology 2008, 375 (1), 223–235. 10.1016/J.VIROL.2008.01.014.

(46) Compton, J. Nucleic Acid Sequence-Based Amplification. Nature 1991, 350 (6313), 91–92. 10.1038/350091A0.

(47) Holmes, E. C.; Twiddy, S. S. The Origin, Emergence and Evolutionary Genetics of Dengue Virus. Infection, Genetics and Evolution 2003, 3 (1), 19–28. 10.1016/S1567-1348(03)00004-2.

(48) Lukashevich, I. S.; Paessler, S.; de la Torre, J. C. Lassa Virus Diversity and Feasibility for Universal Prophylactic Vaccine. F1000Res 2019, 8, 134. 10.12688/f1000research.16989.1.

(49) Chhabra, P.; de Graaf, M.; Parra, G. I.; Chan, M. C. W.; Green, K.; Martella, V.; Wang, Q.; White, P. A.; Katayama, K.; Vennema, H.; Koopmans, M. P. G.; Vinjé, J. Updated Classification of Norovirus Genogroups and Genotypes. J Gen Virol 2019, 100 (10), 1393–1406. 10.1099/JGV.0.001318.

(50) Hemelaar, J. The Origin and Diversity of the HIV-1 Pandemic. Trends Mol Med 2012, 18 (3), 182–192. 10.1016/J.MOLMED.2011.12.001.

(51) Franco, R. A. L.; Brenner, G.; Zocca, V. F. B.; de Paiva, G. B.; Lima, R. N.; Rech, E. L.; Amaral, D. T.; Lins, M. R. C. R.; Pedrolli, D. B. Signal Amplification for Cell-Free Biosensors, an Analog-to-Digital Converter. ACS Synth Biol 2023, 12 (10), 2819–2826. 10.1021/ACSSYNBIO.3C00227/SUPPL_FILE/SB3C00227_SI_004.XLSX.

(52) Hong, F.; Ma, D.; Wu, K.; Mina, L. A.; Luiten, R. C.; Liu, Y.; Yan, H.; Green, A. A. Precise and Programmable Detection of Mutations Using Ultraspecific Riboregulators. Cell 2020, 180 (5), 1018–1032.e16. 10.1016/J.CELL.2020.02.011.

(53) Rozewicki, J.; Li, S.; Amada, K. M.; Standley, D. M.; Katoh, K. MAFFT-DASH: Integrated Protein Sequence and Structural Alignment. Nucleic Acids Res 2019, 47 (W1), W5–W10. 10.1093/NAR/GKZ342.

